# Area under the curve quantification outperforms spectral counting in metaproteomics, but matching between runs is detrimental

**DOI:** 10.64898/2026.04.05.716595

**Authors:** Ayesha Awan, J. Alfredo Blakeley-Ruiz, Manuel Kleiner, Tjorven Hinzke

**Author notes:** Corresponding authors: Ayesha Awan, Manuel Kleiner, Tjorven Hinzke.

## Abstract

Metaproteomics enables the functional characterization of complex microbial communities by detecting and quantifying thousands of proteins. Accurate quantification is essential for deriving meaningful biological inferences from metaproteomes. In mass spectrometry-driven metaproteomics, protein quantification is performed using either MS1-based area under the curve (AUC) or MS2-based peptide spectral counts (SpC). Systematic benchmarking of AUC vs. SpC quantification for metaproteomics, however, is lacking. Additionally, the impact of matching identifications between runs (MBR) on AUC quantification accuracy has not been evaluated. Here, we tested whether AUC or SpC is better suited for quantification in metaproteomics using data-dependent acquisition mass spectrometry data and investigated the impact of MBR on AUC using defined metaproteomics samples with known composition as ground truth. We found that MBR incorrectly transferred peptide and protein identifications and led to a substantial number of falsely identified proteins in samples of known composition. Additionally, when comparing MBR-free AUC data to SpC, AUC data had a wider dynamic range, higher quantitative accuracy, and higher sensitivity for detecting abundance differences. Based on these findings, we recommend MBR-free AUC quantification for metaproteomics analysis, while SpC-based identifications can be used to increase the number of identified proteins to obtain a more comprehensive view of expressed functions.

**Significance of the study:** Although accurate quantification of proteins is crucial for characterizing microbial functions in microbiomes, quantification strategies in metaproteomics are mostly selected based on convention rather than evidence. With this study, we provide an evidence-based framework for the informed selection of quantification strategies in metaproteomics. We highlight the strengths and limitations of the tested approaches with respect to dynamic range, quantitative accuracy, and proteome coverage. Based on our results, we recommend using AUC data without MBR for quantitative metaproteomics.

## Introduction

Metaproteomics enables the simultaneous quantification of thousands to tens of thousands of proteins in complex environmental samples to answer questions ranging from basic biology to environmental and biomedical applications [1–3]. These questions entail, for example, how much each member of the community contributes to the overall proteinaceous biomass [4], how different partners interact in symbioses [5, 6], whether environmental rehabilitation is successful [7–9], how plants interact with their microbiomes [10, 11], and what impact diet has on the composition and function of the gut microbiome [12]. Reliable and accurate protein quantification is essential for inferring biologically meaningful insights from these complex metaproteomics samples. However, there is no ground truth-based consensus on which quantitative approach is best suited for metaproteomic analysis [13]. A well-suited approach would accurately measure the relative abundance of proteins in the context of the sample’s complex matrix and do so across a wide dynamic range, i.e., the range between the lowest and highest quantified values. A high dynamic range is particularly important in metaproteomics because samples containing hundreds to thousands of taxa have proteins that span many orders of magnitude in abundance, and a larger dynamic range enables accurate quantification of small changes in the abundance of biologically relevant proteins and species.

The most commonly used quantification approaches in metaproteomics are label-free shotgun methods adopted from single-organism proteomics, where the sample matrix is a lot less complex than in metaproteomics [14, 15]. Label-free shotgun metaproteomics uses a bottom-up approach where proteins are first digested into peptides, which are then analyzed by liquid chromatography-tandem mass spectrometry (LC-MS/MS) [16, 17]. The mass spectrometry component of LC-MS/MS can then be performed using data-dependent acquisition (DDA) or data-independent acquisition (DIA), which employ different quantification strategies. In this study, we focused on the two quantification strategies used in DDA, because it is presently the more widely used and better characterized approach for metaproteomic analyses.

The first quantification approach we examined uses the MS1 component of MS/MS tandem mass spectrometry. It quantifies peptide abundance by extracting intensity data from peptide ion chromatographic peaks in MS1 and then calculates the maximum intensity or peak area under the curve (AUC). Alternatively, in the second quantification approach, MS2 spectra generated by fragmenting peptides with a collision gas [18] are counted (spectral counts, SpC) and used to determine peptide abundance [15]. However, since the detection and identification of peptides and proteins in both approaches is based on MS2 spectra [18, 19], every identified protein is associated with an SpC value but may not always have a reliable AUC-quantified abundance. This could be especially true for complex metaproteomic samples, where factors such as low-intensity peaks in MS1 could potentially lead to more proteins with an SpC value but not an AUC quantification.

One approach that has been used to help increase the number of AUC quantified proteins is matching between runs, or MBR. MBR transfers peptide identifications from a sample where the peptide was confidently identified using both MS2 and MS1-based data to a different sample that lacks MS2 evidence for that peptide, using mass and retention time alignment. This feature, originally introduced in MaxQuant [20], is also available, e.g., as feature mapper in Proteome Discoverer and as match-between-runs IonQuant [21]. While decreasing data sparsity, MBR can also lead to false-positive identifications even in samples of low complexity through incorrect association of MS2-based evidence from one sample with an MS1-based chromatographic peak in another sample that is similar in retention time and mass [21, 22]. In complex metaproteomes, where mass peak densities are higher in MS1 spectra, these misidentifications might occur more often, with potential impacts on protein identification and quantification. However, the impact of MBR on metaproteomics analysis based on ground-truth data has not been determined. Additionally, there is no comparative evaluation for AUC vs. SpC in metaproteomics beyond the finding that SpC data is better suited for calculating biomass of microbial community members [4]. While comparison of AUC and SpC in single-organism proteomics data found that MS1-based AUC quantification has a higher dynamic range and enables the detection of smaller protein abundance fold changes compared to SpC [23, 24], it remains unclear whether these differences will translate to metaproteomics given the higher complexity of metaproteomics samples.

In this study, we asked whether AUC or SpC data is better suited for metaproteomics quantification in terms of quantitative accuracy and dynamic range, and whether MBR helps to improve metaproteomics AUC data. To answer these questions, we used defined metaproteomic samples and a DDA mass spectrometry approach to generate ground-truth data for evaluating label-free quantification approaches. Using this data, we analyzed how MBR impacts protein identification and quantification in AUC data. We also compared the dynamic range of AUC and SpC data and assessed the accuracy of both methods in terms of how well the abundance values of proteins of specific organisms in the metaproteome corresponded to those measured for the same organism in pure cultures, and how well measured fold changes between defined metaproteomes corresponded to expected changes. Our results show that using MBR leads to the mis-identification and quantification of hundreds of proteins not present in the sample, and that AUC has a wider dynamic range and higher quantitative accuracy than SpC.

## Materials and methods

### Sample and mass spectrometry data generation

We based our analysis of quantitative approaches in metaproteomics on defined metaproteomes. The defined metaproteomes were generated in biological quadruplicates in a previous study [25]. They consisted of known amounts of peptides extracted from pure microbial cultures combined with metaproteome peptide mixtures from either mouse fecal pellets (i.e., a high-complexity matrix) or from sterile corn roots (i.e., a low-complexity matrix, Table 1). In brief, samples were processed using either FASP or S-Trap protocols [26, 27], and mass spectrometry analysis was performed on an LC-MS/MS system consisting of a Dionex UltiMate 3000 RSLCnano (Thermo Fisher Scientific) and an Orbitrap Exploris 480 (Thermo Fisher Scientific).

**Table 1.** Percentage composition of defined metaproteomic samples used in this study. Mix numbers (Mix no.) correspond to those in Hinzke et al, 2025 [25]. HT: high temperature, LT: low temperature, HL: high light, LL: low light, CM: complex medium, MM: minimal medium. M1-M5 were generated using mouse fecal pellet peptides as matrix; M9-M11 were generated using sterile corn root peptides as matrix. In this study, we analyzed the TT and RL data in detail, as TT data spans a wide abundance range (1 to 30% peptide-based abundance) and is not confounded by proteins shared with the metaproteomic matrix, and RL offers the opportunity to study effects of MBR in known presence vs. absence of an organism.

| Mix no. | Matrix metaproteome (mouse fecal pellets/ corn roots) | *Thermus thermophilus* (TT) |  | *Chlamydomonas reinhardtii* |  | *Rhizobium leguminosarum* (RL) |  | *Escherichia coli* |  | *Bacteroides thetaiotaomicron* |
| --- | --- | --- | --- | --- | --- | --- | --- | --- | --- | --- |
|  |  | Cond. 1 (HT) | Cond. 2 (LT) | Cond. 1 (HL) | Cond. 1 (LL) | bv. viciae 3841 | bv. viciae VF39 | Cond. 1 (CM) | Cond. 2 (MM) |  |
| M1 | 65 | 10 |  | 5 |  | 5 |  | 10 |  | 5 |
| M2 | 65 |  | 10 |  | 5 | 5 |  |  | 10 | 5 |
| M3, M9 | 63 | 30 |  | 1 |  |  |  | 1 |  | 5 |
| M4, M10 | 55 | 10 |  | 10 |  |  | 5 |  | 10 | 10 |
| M5, M11 | 78 |  | 1 |  | 5 |  | 10 | 5 |  | 1 |

### Identification and quantification of proteins

For protein identification and quantification, we searched the mass spectra generated against sample-specific databases (see [25] for details) using Proteome Discoverer (PD) v. 2.3 (Thermo Fisher Scientific). Protein identification and quantification were done in PD in two separated steps, a processing step and a consensus step, both of which consist of nodes that do substeps: In the processing step, peptides and proteins were identified by database search in the Sequest HT node, PSM level FDR was calculated in the Percolator node, and MS1 features (for AUC quantification) were identified in the Minora Feature Detector node (Supp. Fig. S1). In the consensus step, protein AUC abundance was calculated in the Feature Mapper and Precursor Ions Quantifier nodes, peptide and protein level FDR was recalculated through a series of validator and filter nodes, and protein groups were formed in the Protein Grouping node (Supp. Fig. 2). In the processing step, MS raw files can either be processed individually (called “As Batch”), generating separate quantification files for each sample (Supp. Fig. 3), or all together generating a multiconsensus, which encompasses a multisample protein quantification matrix. Samples initially run “As Batch” can be reanalyzed as a multiconsensus, generating a multisample protein quantification matrix (Supp. Fig. 4).

For AUC quantification, multiconsensus automatically triggers MBR, regardless of parameter settings and whether samples are processed together from the beginning or are reanalyzed. In order to turn off MBR, samples must be searched “As Batch” and protein quantifications combined into a protein quantification matrix outside of PD using custom code (Supplementary File Code). For spectral counting (SpC) quantification, the Minora Feature Detector, Feature Mapper, and Precursor Ion Quantifier nodes are excluded, and counting peptide spectral matches is used for quantification instead. SpC does not use MBR, so the results are MBR-free regardless of whether a multiconsensus is generated or protein quantities are combined outside of PD. We tested four quantification approaches: AUC without MBR (MBR-free), SpC, AUC with MBR using PD Default Parameters, and AUC with MBR using MaxQuant-derived parameters. For all methods, peptide and protein FDR was controlled at 5%. See Supp. Table 1 for the full parameter list for all four quantification methods. MBR-free AUC samples were processed “As Batch” and combined into a protein quantification matrix (Supp. Table 4) outside of PD using custom code (Supplementary File Code).

The SpC protein quantification matrix was generated in a previous study [25]. For that, samples were initially processed “As Batch” and then reanalyzed with multiconsensus to produce the protein quantification matrix (see InputData folder in 10.5281/zenodo.17880379 and Supp. Tables 3 and 7 [25]). For AUC quantification with MBR, samples were searched together, and a protein quantification matrix was created automatically via multiconsensus. We determined the effect of MBR parameter stringency on protein identification and quantification by using two different Feature Mapper parameter sets (Supp. Table 1) based on PD’s default parameters (Supp. Table 5) and MaxQuant’s (Supp. Table 6) default parameters [20]: (i) Default PD Feature Mapper parameters, i.e., Retention Time (RT) alignment set to True, coarse parameter tuning, 10 min maximum RT shift, Chromatographic Alignment Mass Tolerance of 10 ppm, 0 min RT tolerance, Feature Linking and Mapping Mass Tolerance of 0 ppm, and a minimum signal-to-noise ratio threshold of 5 in the Feature Linking and Mapping section; (ii) Parameters to reflect commonly used parameters in the literature that are often associated with MaxQuant default MBR settings (MBR-MQ) [20, 28], i.e., RT alignment set to True, coarse parameter tuning, 20 min maximum RT shift in the Chromatographic Alignment section and Chromatographic Alignment Mass Tolerance of 5 ppm, 0.7 min RT tolerance, Feature Linking and Mapping Mass Tolerance of 5 ppm and a minimum signal to noise ratio threshold of 5 in the Feature Linking and Mapping section.

### Data analysis and visualization

We used R version 4.2.2 for data analysis. We filtered data to contain only master proteins with a 5% FDR cutoff detected in at least 3 of the 4 replicates in all downstream analyses. AUC quantification-based protein abundance values were normalized using total sum scaling (TSS), and spectral counts were normalized using the normalized spectral abundance factor (NSAF) [29]. To obtain the relative abundances for proteins of a specific species in the metaproteomic samples in SpC data, including *Rhizobium leguminosarum* and *Thermus thermophilus*, we renormalized the NSAF data to sum to 100% at the organism level (orgNSAF%) [4, 30]. To obtain the relative abundance of proteins of a specific species in the metaproteomic samples in AUC data, we renormalized the raw AUC abundances of proteins using TSS on the organism level (orgTSS%). We calculated Spearman’s rank correlations using the sm_statCorr function in the R package smplot2 version 0.2.4 [31, 32]. For all correlation and log2 transformation-based analyses, only proteins with quantification values in both AUC and SpC methods were included in the comparisons. For visualization, we used ggplot2 v. 3.5.2 in R and Adobe Illustrator [33].

## Results

### Defined metaproteomes enable ground truth-based comparisons of AUC vs. SpC quantification performance

We used pure cultures and defined metaproteomes, i.e., complex metaproteomes with known composition, to assess how well protein abundances in metaproteomes represent those in pure cultures for AUC vs. SpC quantification, and how accurately protein fold changes are estimated. Our defined metaproteomes consisted of pure culture peptides from five different organisms mixed into complex metaproteomic matrices of peptides extracted from either mouse fecal pellets (M1-M5) or corn roots (M9, M10, M11). In this study, we analyzed the data of *Thermus thermophilus* (TT) grown under two different conditions (high temperature, TTHT, and low temperature, TTLT) mixed into these defined metaproteomes to see whether metaproteome complexity affected quantification of TT proteins. We chose the TT data in this study, as these span a wide abundance difference from 1-30% peptide-based abundance, and as there is no protein identification overlap between TT and proteins of other organisms in the defined metaproteomes, which would confound quantification. For AUC, we first assessed the effect of MBR to see whether transferring identifications is a suitable strategy to increase the number of AUC-quantified proteins for metaproteomics. For that, we included defined metaproteomes with and without *Rhizobium leguminosarum* (RL) in our analysis to analyze whether MBR leads to misidentifications in a setting of known presence vs. absence of an organism.

### Match between runs leads to false protein identifications in metaproteomic samples

To see the impact of MBR on AUC-based quantification, we compared the abundance of RL proteins in defined metaproteomes with RL (M4, M10: 5% RL, M5, M11: 10% RL) and without RL (M3, M9), when analyzed MBR-free, vs. when analyzed with MBR using PD or MQ settings. We found that, when samples were analyzed MBR-free, as expected, very few proteins (on average 6.2 RL proteins summing to 0.02% in M3 and 16.7 RL proteins summing to 0.07% in M9 across 4 replicates) were quantified for RL (Fig. 1A and B). However, when we used MBR with the default PD settings across all three mixes in each sample matrix (M3, M4, M5; and M9, M10, M11), hundreds of RL proteins were identified and quantified in M3 (576.7 RL proteins, summing to 1% abundance on average across 4 replicates) and M9 (678.7 RL proteins, summing to 1.07% abundance on average across 4 replicates) even though both of these mixes did not contain peptides from RL. While using MQ MBR settings led to fewer (<10%) misidentifications than the PD results described above, the differences between these two settings were minor. That means that regardless of the settings used, misidentifications due to MBR were substantial.

**Figure 1.**
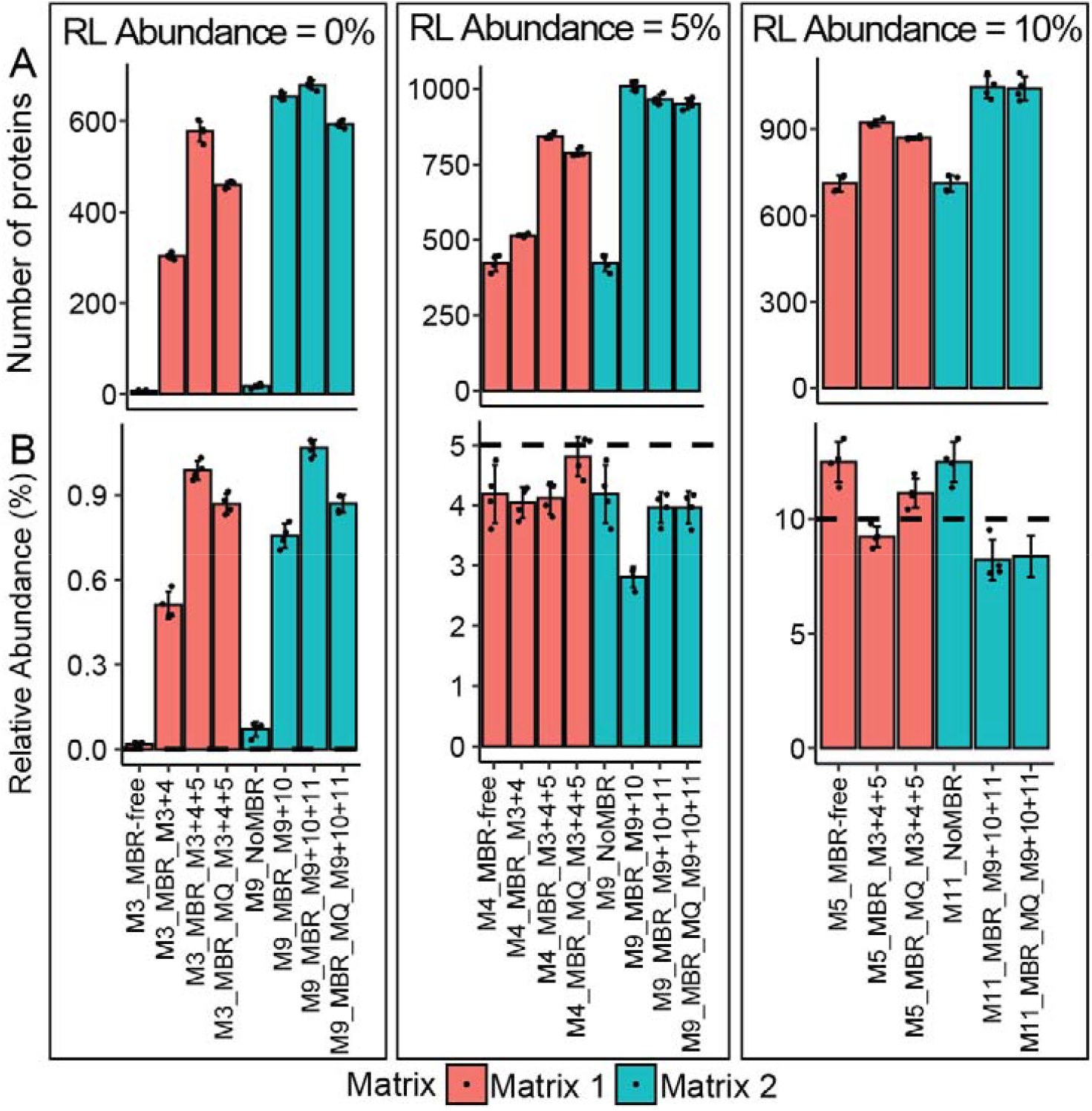
Match between runs (MBR) leads to wrong protein identifications and quantifications. A) Barplot showing the number of *Rhizobium leguminosarum* (RL) proteins identified in mixes with 0% (M3 and M9), 5% (M4 and M10), and 10% (M5 and M11) abundance of RL quantified with and without using MBR. B) Barplot showing the abundance of RL proteins in the same samples as in (A), with and without MBR. To calculate the abundance of RL in the metaproteome in Figure 1B, the relative abundance of all RL proteins in the metaproteome was summed. The dashed line in subplots (B) represents the actual abundance of RL in the defined metaproteome (Table 1). The x-axis labels include the following information separated by underscores: the analyzed sample, the MBR setting, and the additional mixes included in the MBR-based multiconsensus generation. The first term is the analyzed sample or mix number. “MBR-free” indicates that MBR-free data was used, and the final consensus was built using a custom script outside of Proteome Discoverer (PD). “MBR” indicates that MBR was performed using default feature mapping parameters in PD. “MBR_MQ” indicates that MBR was performed with an alternative set of feature-mapping parameters (see Supplementary Table 1). Numbers after “MBR” indicate which additional mixes were included in the analysis. Error bars indicate standard deviations of four biological replicates used for all analyses. Data underlying this figure can be found in Supp. Tables 4-6.

We further investigated to what extent the number of samples combined in searches with MBR impacted these misidentifications. We found that the number of wrong identifications increased with the number of samples and depended on the abundance of RL in the included samples, independent of the complexity of the metaproteome (i.e., results were similar for the highly complex mouse fecal pellet matrix and the less complex sterile corn root matrix): When M3 (0% RL) and M4 (5% RL) were searched together, on average 303.5 RL proteins were identified in M3, with an abundance of 0.5% as opposed to the average of 6.2 RL proteins with a summed relative abundance of 0.02% detected in M3 when it was searched by itself. However, when M5 (10% RL) was used in addition to M3 and M4 for combined searches with MBR, the number and abundance of RL proteins wrongly identified in M3 almost doubled, to 576.7 and 1%, respectively. In M9, which contained no RL and a corn root background matrix rather than the mouse fecal pellet peptides of M3, MBR again produced a higher than expected number and abundance of RL proteins. Specifically, when M9 (0% RL) and M10 (5% RL) were searched together, on average 653.2 RL proteins constituting 0.76% of the total protein abundance were detected in M9 as opposed to the 16.7 RL proteins constituting 0.07% protein abundance in M9 when it was searched by itself. Adding M11 (10% RL) to M9 and M10 for searching increased the number of RL proteins in M9 to 678.7 on average across 4 replicates, with a protein abundance of 1.07%.

### AUC data has a much higher dynamic range and better correlation between pure cultures and metaproteomes than SpC data

After we determined that MBR produces erroneous AUC quantifications, we compared MBR-free AUC vs. SpC quantification in TT pure cultures. Over 80% of TT pure culture proteins were identified and quantified using both AUC and SpC data, while 15-16% of proteins were only identified in SpC data (Fig. 2A). However, most proteins unique to SpC only had 1-3 PSMs associated with them (corresponding to 253 out of 326 or 77.6% of SpC-only proteins in TTLT and 233 out of 302 or 77.2% of SpC-only proteins in TTHT), and the total summed relative abundance of SpC-only proteins was around 3% (Supp. Fig. 5). Additionally in pure cultures, AUC data had a broader dynamic range than SpC data (Fig. 2B). SpC abundance values were more compressed compared to AUC abundance values and in some cases, the data density distribution also indicated that SpC data had a multimodal distribution for low-abundant proteins.

**Figure 2.**
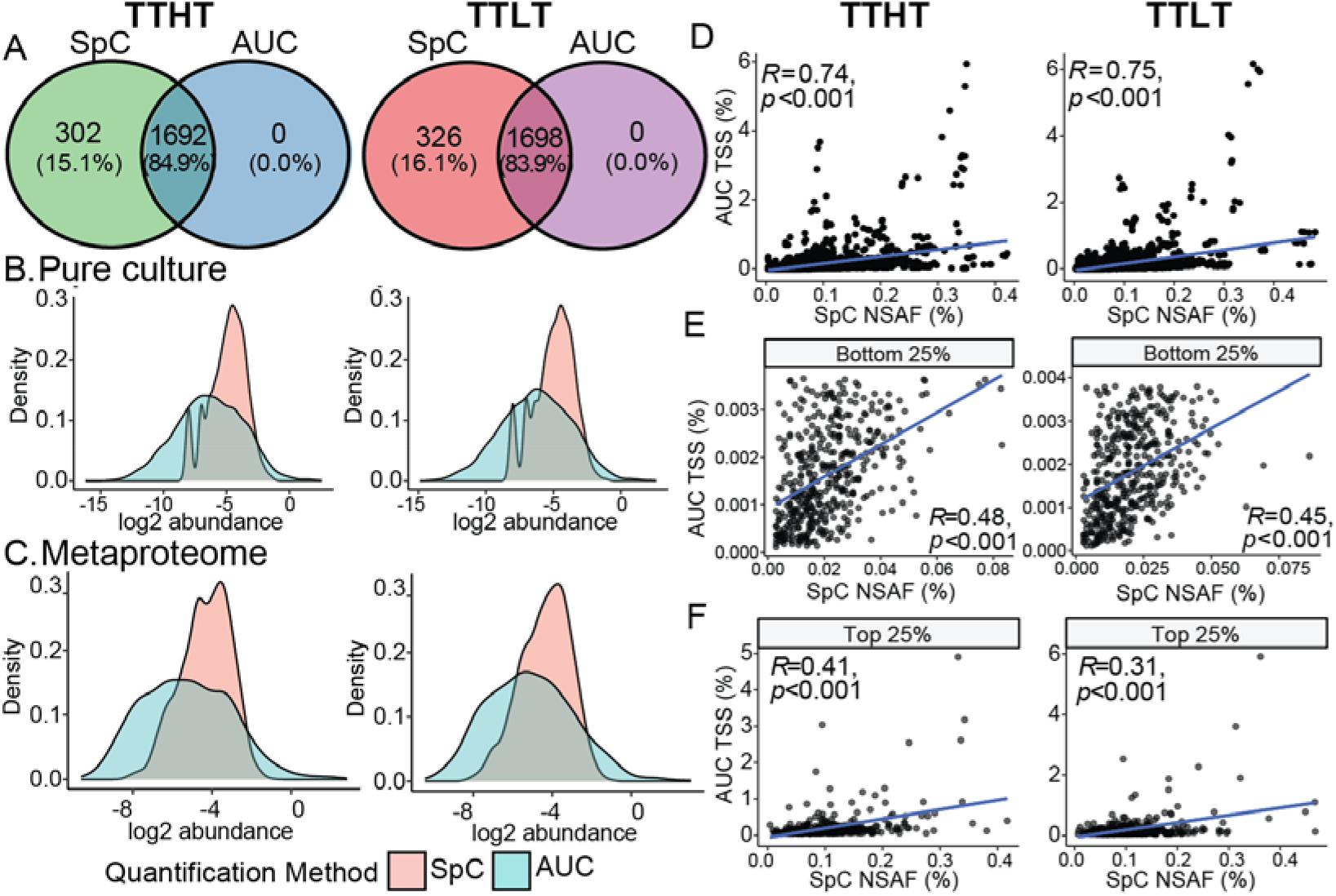
SpC and AUC have highly correlated protein quantification in pure culture proteomes, but AUC has a higher dynamic range in both proteomic and metaproteomic samples. A) Venn diagram showing the number of proteins quantified using SpC and AUC in two different *Thermus thermophilus* (TT) pure culture proteomes (TTHT: TT cultivated under high temperature; TTLT: TT cultivated under low temperature). B) Density plot showing the dynamic range of log2-transformed TT protein abundances quantified using SpC and AUC in pure culture proteomes and C) in defined metaproteomes (M1 and M2) for TTHT and TTLT. D) Spearman’s rank correlation of TT protein abundances quantified using SpC and AUC in TTHT and TTLT pure culture proteomes. E) Correlations between the lowest abundant bottom 25% quartile proteins and F) the highest abundant top 25% quartile proteins are shown separately. AUC abundance values were used to filter for the bottom and top 25% abundant proteins. Pure culture data shown in this figure can be found in Supp. Tables 2 and 3. Metaproteomics data shown in this figure can be found in Supp. Tables 4 and 7.

Using Spearman’s rank based correlation, we assessed concordance between AUC and SpC in isolate proteomes and found that protein abundance ranks were consistent between AUC and SpC quantification data, as shown by Spearman’s rank correlations for both TTHT (R = 0.74; p < 0.001) and TTLT (R = 0.75; p < 0.001) (Fig. 2D). However, when analyzing the upper and lower protein abundance quartiles separately, we found that the correlation between AUC and SpC quantification was weaker for the highest-abundant proteins (TTHT: R = 0.41, p < 0.001; TTLT: R = 0.31, p < 0.001) and the lowest-abundance proteins (TTHT: R = 0.48, p < 0.001; TTLT: R = 0.45, p < 0.001) (Fig. 2E and F). Proteins such as the S-layer protein (Q72HF5) and the periplasmic dipeptide binding protein of an ABC transporter (Q72I61) were much more abundant in the AUC data (∼ 5%, 3%) than in the SpC data (∼ 0.3%, 0.1%).

Consistent with our findings in the pure cultures, fewer proteins were quantified in metaproteomic samples using AUC than were detected in SpC in both TTHT (985 using AUC versus 1,453 using SpC) and TTLT (931 using AUC versus 1,465 using SpC) (Fig. 3A, Supp. Fig. 6A). Similarly to the pure culture data, in defined metaproteomes most TT proteins detected only with SpC were very low-abundant with 1-3 PSMs (460 out of 536 or 85.8% of SpC-only proteins in TTLT and 395 out of 468 or 84.4% of SpC-only proteins in TTHT) (Supp. Fig. 6B), while their summed-up relative abundance was between 12 and 17% of all TT proteins (Supp. Fig. 6C). When considering all proteins in the defined metaproteomes, 80% of the SpC-only proteins had just 1 PSM but together made up almost a third of the total relative protein abundance (Supp. Fig. 7).

**Figure 3.**
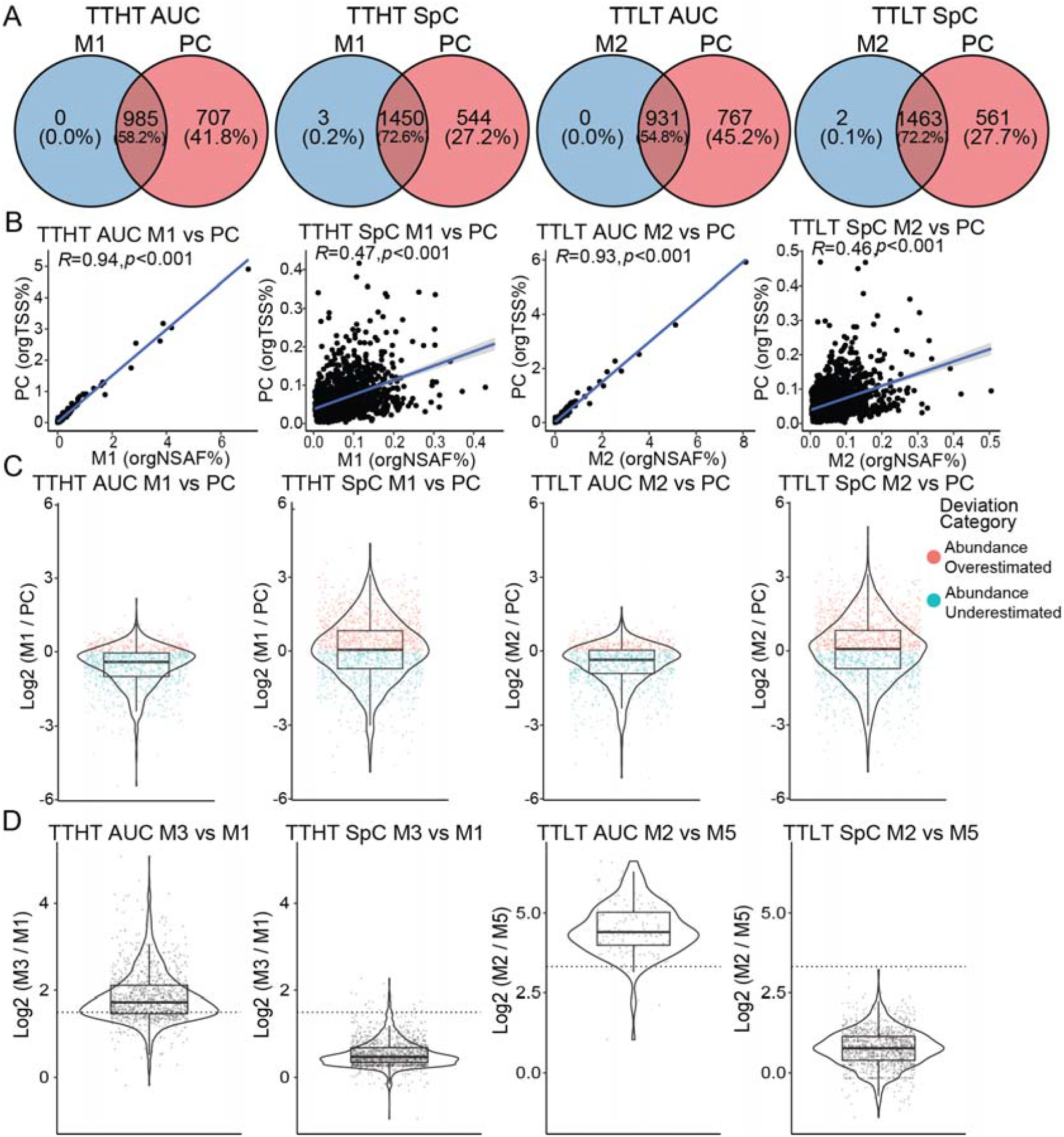
In metaproteomic samples, AUC quantification is more accurate than SpC. A) Number of TT proteins quantified using AUC and SpC in defined metaproteome samples (See Table 1 for composition of mixes). B) Spearman’s rank correlation between the abundance of TT proteins in pure cultures (PC) and defined metaproteomes (M1 and M2) for both AUC (orgTSS%) and SpC (orgNSAF%) quantified relative protein abundances. C) Violin plot with boxplot overlay showing the log2 fold deviation of protein abundances in metaproteomes (M1 and M2) compared to their abundance in pure culture (PC). D) Violin plot with boxplot overlay showing the log2 fold change in protein abundance in defined metaproteomic samples containing different abundances of TT quantified using AUC and SpC. TTHT M3, which contained 30% TT, was compared to M1, which contained 10% TT, and TTLT M2, containing 10% TT, was compared to M5, which contained 1% TT. The dotted line shows the expected log2 fold change. The composition of all mixes is shown in Table 1. The pure culture data used are available in Supp. Tables 2 and 3, and the metaproteomics data used are available in Supp. Tables 4 and 7.

In addition to many proteins detected with only 1 PSM, SpC-based quantification had much lower dynamic range than AUC data and showed lower correlation between pure cultures and metaproteomes than AUC-based quantification (Fig. 2C and 3B). This higher correlation between pure cultures and defined metaproteomes in AUC-based protein abundances was seen for both TTHT (AUC: ρ = 0.94, p < 0.001; SpC: ρ = 0.47, p < 0.001) and TTLT data (AUC: ρ = 0.93, p < 0.001; SpC: ρ = 0.46, p < 0.001; Spearman’s rank based correlations; Fig. 3B). Similar to what was observed in the pure culture proteomes, proteins such as the S-layer protein and dipeptide binding protein were more abundant in the AUC data (7% and 4%) than in the SpC data (0.2% and 0.3%).

### AUC has higher quantitative accuracy in complex metaproteomes compared to SpC

To assess the accuracy of AUC and SpC quantification in metaproteomics samples, we calculated the deviations of TT protein abundances in defined metaproteomes compared to their abundance in pure cultures using both AUC and SpC. We found that with AUC, overestimation of protein abundance occurred less frequently and was less pronounced than with SpC (Fig. 3C). Based on this, we calculated TT protein abundance fold changes across different defined metaproteomes (see Table 1) using both AUC and SpC data and found that AUC more accurately captured differences in protein abundance. When comparing Matrix 1 samples with 10% abundance of TT (M1) to mixes with 30% TT abundance (M3), AUC more closely reflected the expected log2 fold change (∼1.5) than SpC (Fig. 3D). Similarly, when mixes with 10% TT abundance (M2) were compared to mixes with 1% abundance (M5), AUC more accurately reflected the expected log2 fold change (∼3.3) compared to SpC. In contrast, SpC led to consistently and substantially underestimated abundance differences in both cases. Across all comparisons, SpC more frequently underestimated log2 fold changes, while AUC more frequently overestimated fold changes.

## Discussion

In this study, we used defined metaproteomics samples to compare the two main approaches used to quantify relative protein abundance in metaproteomics, i.e., MS1-based AUC and MS2-based SpC, and elucidated how MBR affects AUC quantification. We found that when MBR is used for AUC data, hundreds of proteins are wrongly identified and quantified. While the summed relative protein abundance of known misidentified proteins was low, only about 1% of the total proteome, this can still distort the abundance of other proteins in the metaproteome and consequently the biological conclusions drawn. For example, low-biomass members of a community can be disproportionately important for community functioning [34], and the false detection of the presence of an organism and the false detection of functions are of substantial concern for understanding physiological phenotypes of communities.

The lower number of proteins associated with an AUC quantification than with an SpC quantification could be caused by the fact that MS1-based quantification requires a well-defined chromatographic peak with a high enough signal-to-noise ratio to be considered. Additionally, some parameters used for AUC quantification, such as the exclusive use of high-confidence PSMs for chromatographic feature detection (Minora Feature Detector), reduce the amount of MS2 evidence (PSM) considered for assigning identity to chromatographic peaks. MS2-based SpC quantification, on the other hand, relies only on peptide identification with MS/MS spectra, hence allowing peptides with poor precursor chromatographic peak resolution to still contribute to the final protein quantification. Additionally, medium confidence PSMs are also considered in the SpC count. Therefore, some of the differences in the number of proteins detected and quantified with AUC vs. SpC are due to specific parameters in our AUC and SpC search and quantification methods, and adjusting these parameters might change the outcomes in terms of the number of proteins.

The higher dynamic range of AUC compared to SpC for metaproteomic quantification is consistent with results from single-organism proteomic data [23, 35]. In contrast to SpC, AUC is not impacted by dynamic exclusion in DDA, which limits the number of times an abundant peptide ion is resampled for fragmentation and MS/MS data acquisition. Dynamic exclusion essentially limits the maximum PSM count that can be obtained for a single peptide. AUC also does not have a fixed lower abundance limit, which is numerically fixed to 1 for SpC. This difference in dynamic range also explains the discrepancy we observed in protein abundance between AUC and SpC for highly abundant proteins. These differences are highly relevant for biological interpretation. For example, the *T. thermophilus* S-layer protein, which is expected to be highly abundant in the proteome and account for roughly 5-15% of cellular protein [36–38], was much more abundant and consequently better aligned to these expectations according to AUC in both the pure culture proteome (up to ∼5%) and the metaproteome (up to ∼7%) as compared to SpC (up to 0.3% in both proteome and metaproteome). This indicates that AUC can more accurately quantify relevant molecular phenotypes compared to SpC.

Likewise, the discrepancy between AUC and SpC for low-abundant proteins is expected due to the numerical properties of SpC data, which are discrete count data. AUC based abundance values, in contrast, are not discrete counts, but are continuous data that provide much finer resolution for abundance differences for low-abundance proteins. These properties of high dynamic range and continuous data also translate into higher accuracy of AUC data compared to SpC, as shown by the higher correlation between pure culture and metaproteomics abundance data in AUC quantifications. Higher correlation in the relative abundance of species-specific proteins in the context of a pure culture and a complex metaproteome for the same species reflects that AUC data accurately captures protein abundance regardless of the level of complexity of the sample matrix, which is essential for deriving meaningful biological conclusions in metaproteomic samples. AUC also performed better than SpC at capturing fold-changes between the same proteins in different metaproteomics samples, likely due to its wider dynamic range and greater accuracy, which enables it to better detect abundance differences in proteins at both the high and low abundance extremes.

Based on our results, we recommend that if the goal of a metaproteomics study is to accurately quantify protein abundances and protein fold changes that reflect biologically relevant metaproteomic shifts, AUC quantification without MBR should be used. We would like to stress that accurate protein quantification is distinct from protein detection or identification, or the testing of statistical differences in protein abundance between conditions. Proteins identified using SpC that lack AUC quantification can be used in addition to those with AUC quantification to assess pathway completeness, and to sum up relative amounts of functional categories, as well as to perform presence-absence analysis, and to assess qualitative differences [25]. Approaches to combine AUC and SpC data have been proposed [39, 40], but are beyond the scope of this study.

Our study does not investigate pending quantification questions in data-independent acquisition (DIA)-based workflows, which are being increasingly employed for metaproteomics studies [41, 42]. However, DIA has several additional open questions pertaining to database strategies, statistical evaluation, and general data reliability that require further ground truth-based evaluation and validation [42–44]. At the same time, our results hold promise for the usability of purely intensity-based quantification in DIA [45]. We plan on extending the framework presented here to DIA data in the future.

## Supporting information

Supplemental Results

## Data availability

Proteomics data used in this study are available at the ProteomeXchange Consortium (http://proteomecentral.proteomexchange.org) via the PRIDE repository with the data set identifier PXD045390 (Reviewer access: Username; Password: FwNk89lz).

## Supplementary materials

Supplementary Table 1: Table describing all parameters used for each of the 4 quantification methods. The parameters are almost identical, and the differences are in how the search was performed together or separately, and changes to the Minora Feature Detector and Feature Mapper nodes (highlighted in blue).

Supplementary Table 2: MBR-free AUC quantified protein abundances in *T. thermophilus* pure cultures

Supplementary Table 3: SpC quantified protein abundances in *T. thermophilus* pure cultures

Supplementary Table 4: MBR-free AUC quantified protein abundances in Matrix 1 Mixes M1-M5 and in Matrix 2 Mixes M9-M11

Supplementary Table 5: AUC quantified protein abundances in Matrix 1 Mixes M3-M5 and in Matrix 2 Mixes M9-M11 with MBR (PD default parameters)

Supplementary Table 6: AUC quantified protein abundances in Matrix 1 Mixes M3-M5 and in Matrix 2 Mixes M9-M11 with MBR (MaxQuant-derived parameters)

Supplementary Table 7: SpC quantified protein abundances in Matrix 1, Mix 1 and 2

Supplementary Code: Annotated Python script used to build consensus protein tables outside of PD

## Declaration of interests

The authors declare that they have no competing interests.

## Author contributions

AA: Data analysis, figure generation, writing and editing the manuscript

JAB: Data analysis, writing and editing the manuscript

MK: Funding, conceptualization of the study, experimental design, data analysis, writing and editing the manuscript

TH: Funding, conceptualization of the study, experimental design, data analysis, writing and editing the manuscript

## Acknowledgements

All LC-MS/MS measurements were made in the Molecular Education, Technology, and Research Innovation Center (METRIC) at North Carolina State University. The authors would like to thank Clara Tang for her assistance in formulating the early stages of this project. This research was funded by the Deutsche Forschungsgemeinschaft (DFG, German Research Foundation) – Project-ID 531801029 – TRR 410 “WETSCAPES2.0” and Project-ID 522593772, the U.S. Department of Agriculture National Institute of Food and Agriculture under award No. 2022-67013-36672, the US National Science Foundation under award number IOS #2426305, and the National Institute Of General Medical Sciences of the National Institutes of Health under Award Number R35GM138362. The content is solely the responsibility of the authors and does not necessarily represent the official views of the National Institutes of Health or the National Science Foundation.

## Notes

### Competing Interest Statement

The authors have declared no competing interest.

### Summary of Updates

Results and supplementary results updated to include more analyses.

