## Supplemental Results for "Area under the curve quantification outperforms spectral counting in metaproteomics, but matching between runs is detrimental"

Corresponding authors:

### Supplementary Methods

We searched raw mass spectrometry data against custom databases using ProteomeDiscoverer v. 2.3 (Thermo Fisher Scientific, see main manuscript for details). For screenshots of the used workflow, refer to Supp. Figs 1-4.

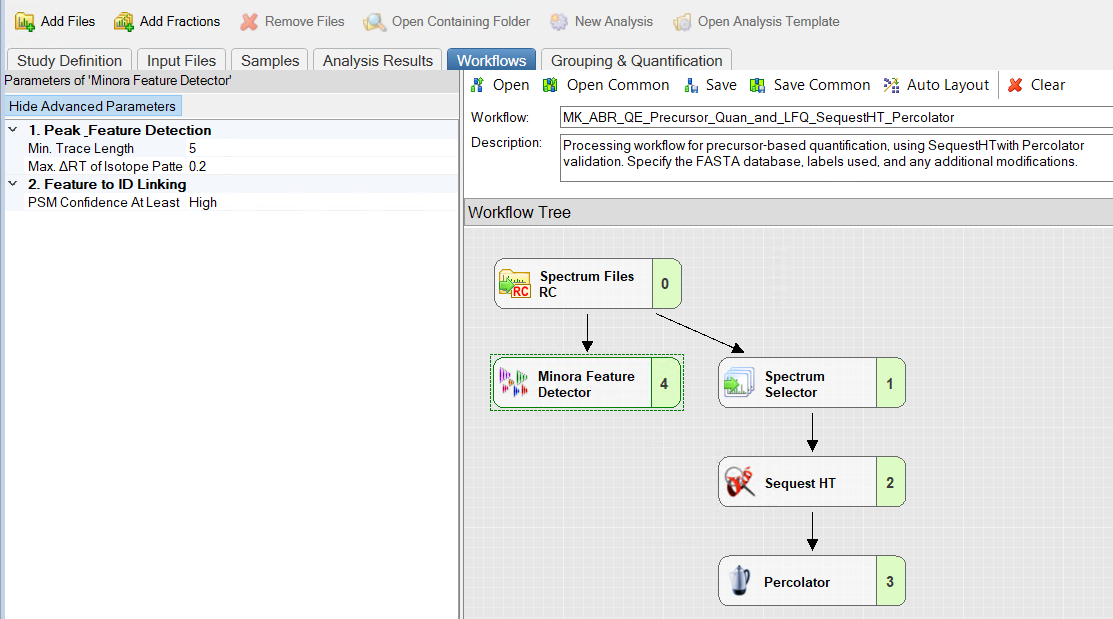

**Supplementary Figure 1**: Screenshot of the ProteomeDiscoverer processing workflow as used in this study. The Minora Feature Detector is highlighted. The Minora Feature Detector node was excluded for SpC.

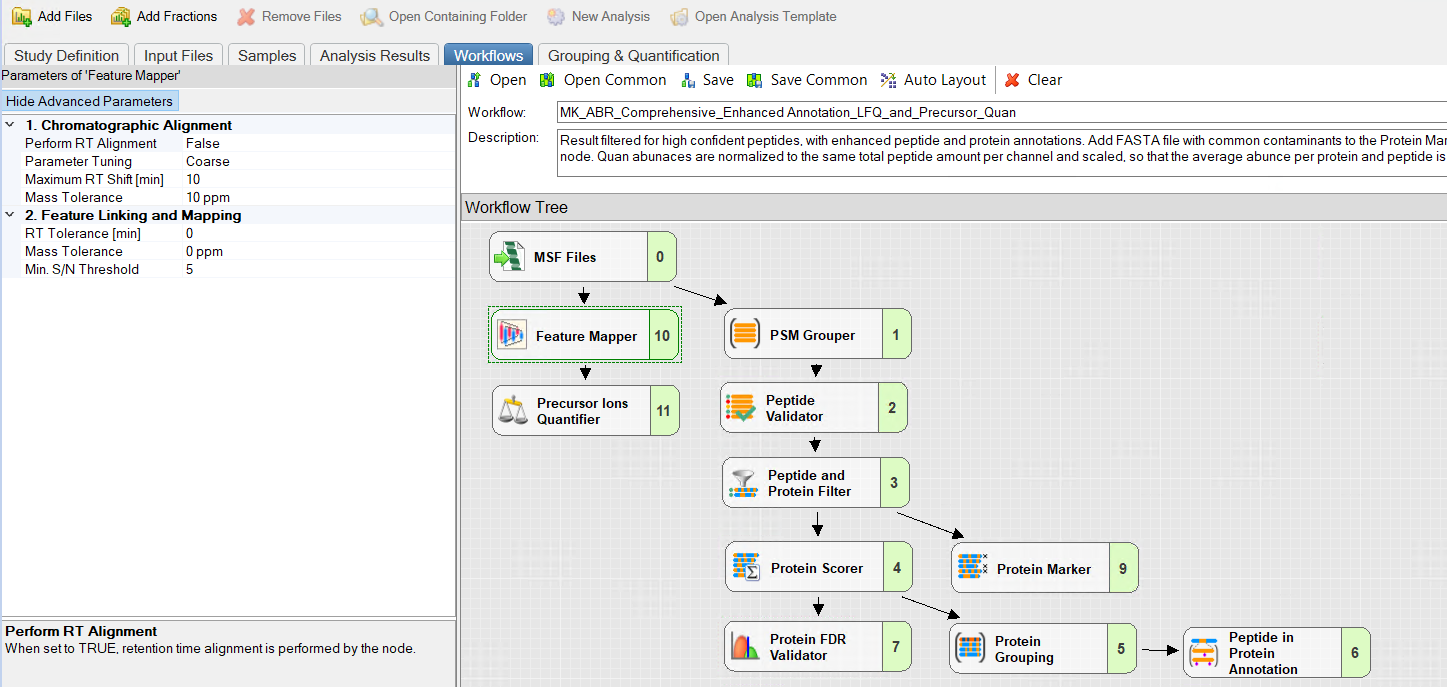

**Supplementary Figure 2**: Screenshot of the ProteomeDiscoverer Consensus Workflow as used in this study. The Feature Mapper Node is highlighted. The following parameters were changed here between the PD default parameters and the MaxQuant simulating parameters: Chromatographic Alignment: Maximum RT Shift [min], Chromatographic Alignment: Mass Tolerance, Feature Linking and Mapping: RT Tolerance [min], and Feature Linking and Mapping: Mass Tolerance; see Supp. Table 1 for the respective parameters. Feature Mapper and Precursor Ion Quantifier are excluded in SpC search.

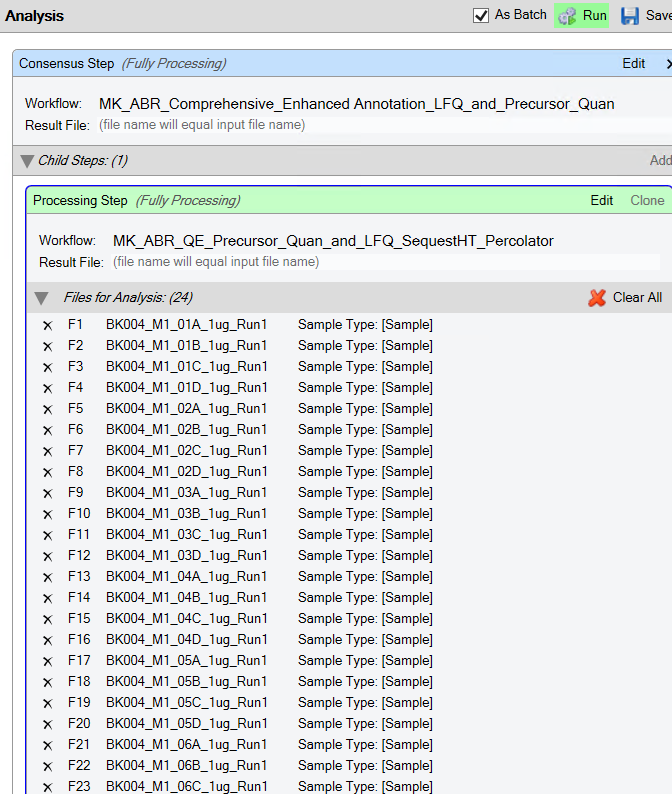

**Supplementary Figure 3**: Screenshot of the ProteomeDiscoverer settings for starting a (meta)proteomics raw file analysis for protein identification and quantification. Raw files are placed under the processing step. If the “As Batch” button is activated, the raw files are searched separately, producing separate output files that can be combined into a single file outside of ProteomeDiscoverer, e.g., using the script used in this study, which we provide as Supplementary Code. If the “As Batch” button is not activated, then the raw files are searched together, resulting in a single consensus file output.

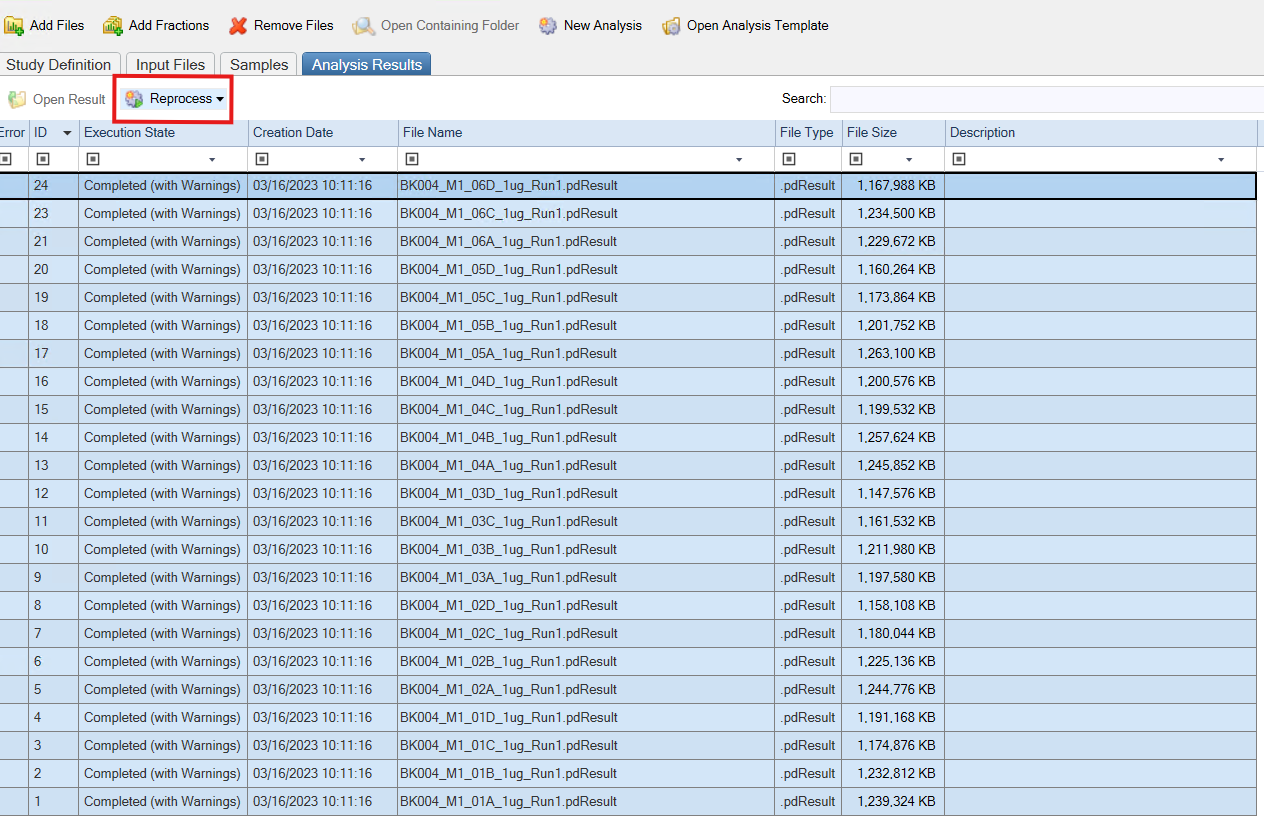

**Supplementary Figure 4**: Screenshot of reprocessing files analyzed “As Batch” to create a new multiconsensus. We only did this step for the SpC data.

### Supplementary Results

In pure cultures as well as in defined metaproteomes, a substantial number (300-600) of TT proteins were only detected via SpC, but not quantified via AUC. The majority of these SpC-only proteins were low-abundant with 1-3 PSMs (Supp. Figs 5 and 6). As *Thermus thermophilus* (TT) proteins detected only via SpC in metaproteomes were also detected in pure culture using SpC, these proteins can be confidently designated as truly being present and not misidentifications. Summed together, low-abundant, SpC-only proteins from all taxa in M1 also made up a considerable relative amount of the metaproteome (Supp. Fig 7). While these SpC-only proteins thus need to be considered in a qualitative functional analysis, and can be used for statistical testing [[1]](https://www.zotero.org/google-docs/?ZTKuwz), they lack the accuracy of AUC data for quantitative analyses.

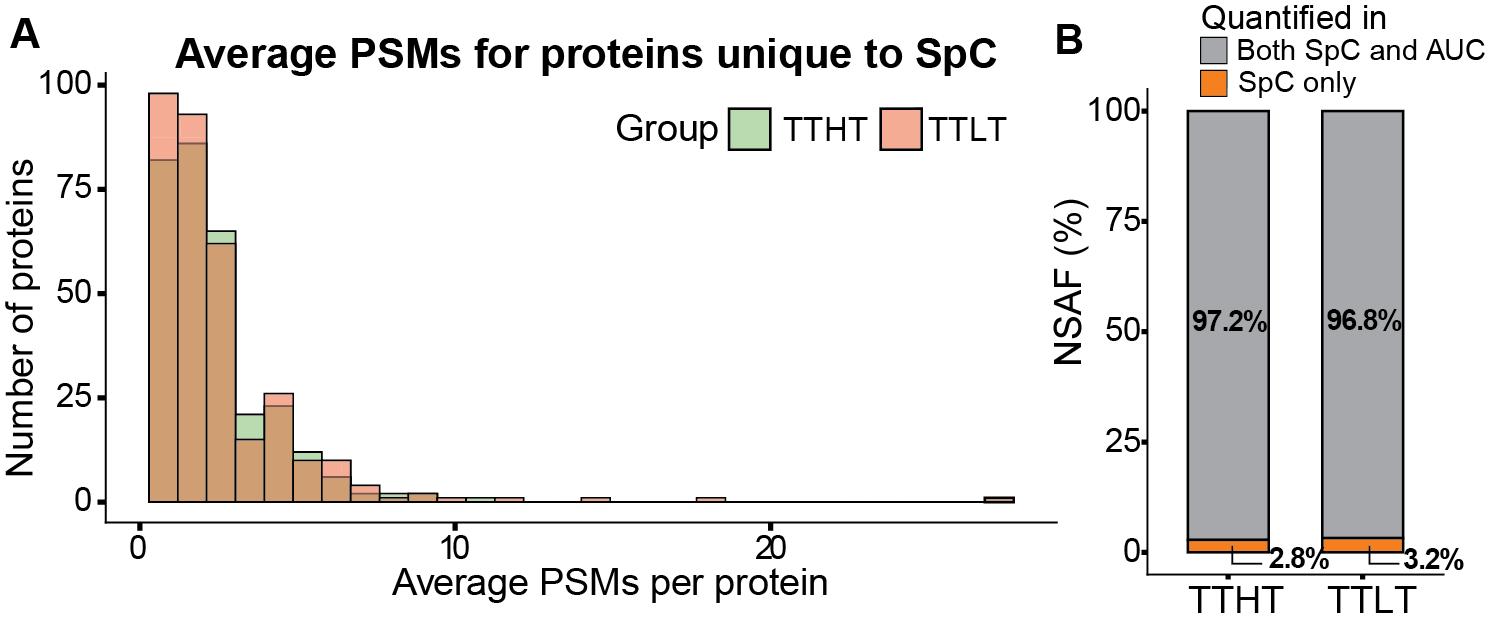

**Supplementary Figure 5:** Proteins detected only via SpC, but not quantified via AUC, in pure cultures are mostly very low-abundant. A) Average PSMs per protein across replicates detected only with SpC data, but not quantified with AUC data in *Thermus thermophilus* (TT) pure cultures grown under high (HT) and low (LT) temperature. B) Relative abundance of TT proteins detected only with SpC data, but not quantified with AUC data.

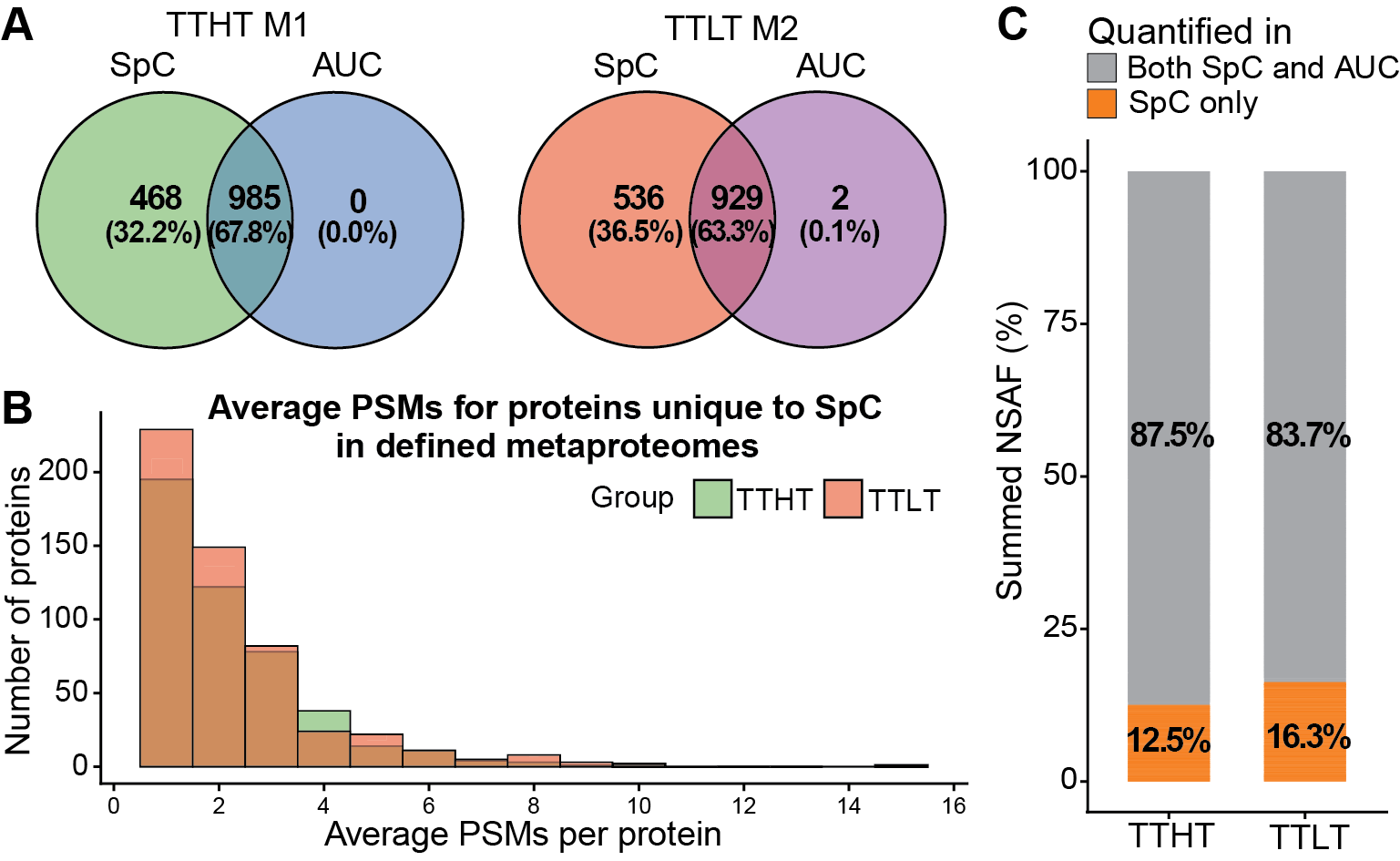

**Supplementary Figure 6**: About one third of proteins are detected only via SpC, but not quantified via AUC in defined metaproteomes, and are mostly low-abundant. A) Venn diagram showing the number of proteins quantified using SpC and AUC in *Thermus thermophilus* (TTHT: TT cultivated at high temperature; TTLT: TT cultivated in low temperature) in defined metaproteomes. B) Average PSMs per protein detected only with SpC data, but not quantified with AUC data in TT grown under high (HT) and low (LT) temperature in defined metaproteomes. C) Relative abundance of TT proteins detected only with SpC data, but not quantified with AUC data in defined metaproteomes.

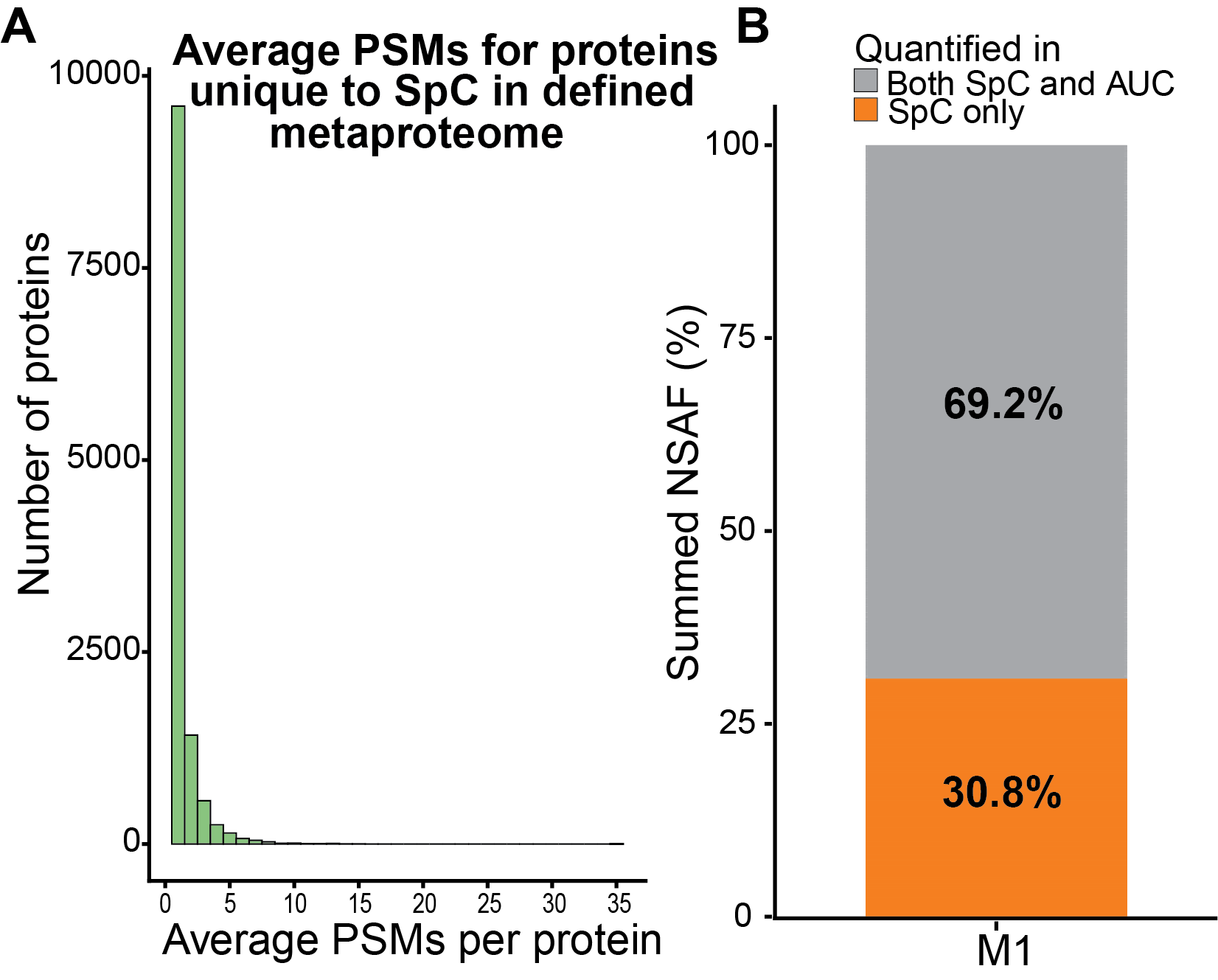

**Supplementary Figure 7**: While most proteins only detected via SpC, but not quantified via AUC, in defined metaproteomes are very low-abundant, they add up to a substantial part of the overall metaproteome. A) Average PSMs per protein detected only with SpC data, but not quantified with AUC data in defined metaproteome M1. B) Relative abundance of proteins detected only with SpC data, but not quantified with AUC data.
