## Supplementary Methods Code for "Area under the curve quantification outperforms spectral counting in metaproteomics, but matching between runs is detrimental"

#MergePDProteinResults.py
#Writen J. A. Blakeley-Ruiz
#Purpose: This code will merge multiple proteome discoverer v. 2.3 (PD)
#protein results files into a single matrix with protein id 
#in the first column and AUC abundances by sample in the second column
#Default filter is 5% FDR and master protein classification (Filtering criterea can be adjusted in rows 72 and 73)
#Usage: This code should work with any version of python 3, command: python3 SupplementaryMethodsCode.py PathToFolder Output


#Required python modules
import os
import sys
import pandas as pd
import csv

#Make a list of all files
path = sys.argv[1] #Taken from the PathToFolder Slot in the command to run the code.
ProteinlistOfFile = [] #List that holds file path for all protein tables
ProteinFileName = [] # List that holds file names for all protein tables
PSMlistOfFile = [] # List that holds file path for all PSM tables
PSMFileName = [] # List that holds file names for all PSM tables
MS2listOfFile = [] # List that holds file path for all MS2 tables
MS2FileName = [] # List that holds names path for all MS2 tables

#Pulls file names from folder that contains PD results files. 
for (dirpath, dirnames, filenames) in os.walk(path): #loops through folder and pulls all the file paths and names
    for filename in filenames:
        tmp = os.sep.join([dirpath, filename])
        
        #Pulls protein table paths and file names
        if "_Proteins" in filename:
            ProteinlistOfFile.append(tmp)
            ProteinFileName.append(filename)
        #Pulls PSM table paths and file names
        elif "_PSMs" in filename:
            PSMlistOfFile.append(tmp)
            PSMFileName.append(filename)
        #Pulls MS2 table paths and file names
        elif "_MSMSSpectrumInfo" in filename:
            MS2listOfFile.append(tmp)
            MS2FileName.append(filename)

#Matrices That Will Hold Data
Data = {}
ProteinAA = {}
ProteinUniquePeptides = {}

#Merge Protein.txt files into one Matrix
for j in range(0,len(ProteinlistOfFile)): #Loops through all the file paths in the ProteinlistOfFile list
    print(ProteinlistOfFile[j]) #Reports all the file paths loops through and can be used to trouble shoot if there is an error. The last file path to appear is the last file attempted to be opened. 
    with open(ProteinlistOfFile[j]) as f: #Opens next file in the list
        file = csv.reader(f, delimiter='\t') #module that reads the protein.txt file
        header = next(file) #puts the first line of the protein.txt file in a list called header. 
        #Data[ProteinFileName[j]] = {}
        for i in range(0,len(header)): #Checks the header list for specific columns in the protein.txt and puts them in a variable, works for both R friendly and default .txt export setting for PD.
            
            #Collects index for protein level FDR
            if header[i] == "Exp q-value Combined":
                Q = i
            elif header[i] == "Exp. q-value: Combined":
                Q = i
            #Collects index for protein unique peptides
            elif header[i] == "Number of Protein Unique Peptides":
                UniquePep = i
            elif header[i] == "# Protein Unique Peptides":
                UniquePep = i
            #Collects index for protein ID
            elif header[i] == "Accession":
                Accession = i
            elif header[i] == "Number of PSMs":
                PSM = i
            elif header[i] == "# PSMs":
                PSM = i
            #Collects index for master protein column
            elif header[i] == "Master":
                Master = i
            #Colects index for # of amino acids column
            elif header[i] == "Number of AAs":
            	AA = i
            elif header[i] == "# AAs":
                AA = i   
            #Collects index for protein description column
            elif header[i] == "Description":
                Descript = i
            #Collects index for sequence column. 
            elif header[i] == "Sequence":
                Seq = i
            #Collects index AUC abundance column 
            elif "Abundance:" in header [i]:
                Abundance = i
        for row in file: #Set filtering critera
            if float(row[Q]) <= 0.05: #This number sets the protein level FDR control. 0.05 = 5% FDR lower numbers decrease FDR and higher numbers in crease FDR
                if row[Master] == "IsMasterProtein": # This step limits the identified proteins to those that have been determined to be present in the sample by parsimony rules of unique and shared peptides. Remove if not desired and can be changed to unique peptides by changing to "if row[UniquePep] > 0"
                    
                    #If protein already in Data dictionary then only the abundance is added to the dictionary
                    if row[Accession] in Data.keys():
                        Data[row[Accession]][ProteinFileName[j]] = row[Abundance] #Change "Abundance" to "PSM" to quantify by PSM instead.
                        ProteinUniquePeptides[row[Accession]][ProteinFileName[j]] = row[UniquePep] # Counts the number of unique peptides detected for the protein in sample
                    #If this is the first instance of the protein's detection than the proteins abundance dictionary will be created. 
                    else:
                        Data[row[Accession]] = {}
                        Data[row[Accession]]["AALength"] = row[AA] #Records amino acid length of protein
                        Data[row[Accession]]["Description"] = row[Descript] # Records protein description if recorded by PD. 
                        Data[row[Accession]]["Sequence"] = row[Seq] # Records sequence of protein. IMPORTANT: for very large proteins like Titin this can break the output whan opened in Excel. Comment out this line to avoid.
                        #Data[row[Accession]][ProteinFileName[j]]= row[PSM] #Remove this comment to quantify by PSM instead
                        Data[row[Accession]][ProteinFileName[j]] = row[Abundance] #Quantifies by AUC comment this line and remove comment in previous line to quantify by PSM instead
                        ProteinUniquePeptides[row[Accession]] = {} 
                        ProteinUniquePeptides[row[Accession]][ProteinFileName[j]] = row[UniquePep] # Counts the number of unique peptides detected for the protein in sample


                    

#Save table
df = pd.DataFrame.from_dict(Data, orient="index")
df.to_csv(sys.argv[2],sep="\t")

#df = pd.DataFrame.from_dict(ProteinUniquePeptides, orient="index")
#df.to_csv("ListOfProteinUniquePeptides.csv",sep="\t")
#tmp = "UniquePep_"+ sys.argv[2]
#df.to_csv(tmp,sep="\t")
